# Development of Permanent Artificial Bowel Replacement Substrates

**DOI:** 10.1101/2021.10.23.465560

**Authors:** Kantida Koysombat, Atikah Haneef

## Abstract

Short bowel syndrome (SDS) is a malabsorption disorder caused by loss of function of the small intestine, either by trauma or innately. Current treatment options include parenteral nutrition (PN) or allograft transplants. Long term PN dependence can lead to complications due to line infections and toxicity from the formula itself. A lack of healthy donors results in long waiting lists and high mortality rates. With allograft transplants, long-term graft and patient survival rates are poor (48% and 39% respectively at 5 years); graft loss occurs due to rejection (48%), thrombosis (28%), sepsis (12%); main causes of death are due to bacterial infection (94%) and rejection. Costs associated with PN annually per patient approximate to £40,000, whereas one allograft procedure costs approximately £80,000; not including intervention due to complications.

Interest in developing an off-the-shelf bioengineered alternative have been expressed. Autologous transplants could be a more beneficial route to improving survival rates, enabling the transplant of patients’ healthy cells back to them. We describe here the development of a synthetic poly(ethylene terephthalate) scaffold using electrospinning, which showed excellent physical and chemical characteristics; high surface area:volume ratio, high mechanical strength, high porosity, and the ability to be chemically/physically functionalised without losing integrity in structure and physical properties. The cost of electrospinning is far lower in comparison to the current available treatment options, potentially providing a stable, off-the-shelf, ready-to-culture product as the need arises for applications in tissue engineered small intestine (TESI), or 3D models for small bowel research.

## Introduction

Short bowel syndrome is a malabsorption disorder which occurs due to the physical loss or the loss of function of a portion of the small intestine, either at birth or after surgery (Tappenden 2014). Current treatment options include parenteral nutrition (PN) or allograft transplants. Long term PN dependence can lead to complications due to line infections and toxicity from the formula itself. A lack of healthy donors results in long waiting lists and high mortality rates worldwide (Schalamon, Johannes et al. 2004). With allograft transplants, long-term graft and patient survival rates are poor (48% and 39% respectively at 5 years) (SJ Middleton et al 2004); graft loss occurs due to rejection (48%), thrombosis (28%), sepsis (12%); main causes of death are due to bacterial infection (94%) and rejection. Costs associated with PN annually per patient approximate to £40,000, whereas one allograft procedure costs approximately £80,000; not including intervention due to complications.(Bharadwaj et al 2017)

Therefore, development of a synthetic bowel substrate which can be readily available as an off-the-shelf product has been proposed. The principal of bowel tissue engineering consists of seeding autologous cells onto a synthetic scaffold, followed by in vitro tissue maturation and ultimately implanting the construct into the host environment (Lanza, Langer et al. 2007). The specific physical properties of bowel substrates that are crucial for the success of this approach are biocompatibility, regulated permeability and suitable mechanical properties allowing the material to withstand peristaltic actions. The scaffold should aim to mimic in vivo microenvironments of the targeted extracellular matrix (ECM) by replicating biochemical, biomechanical and topographical cues to regulate specific cellular response.

Basement membrane (BM) is a form of ECM that provides anchorage to cells and separates epithelium from underlying lamina propria (Vancamelbeke and Vermeire, 2017). Naturally, it is composed of 4 main proteins, including laminin, type IV collagen, the glycoprotein nidrogen and the heparan sulfate proteoglycan perlecan and carbohydrate polymers such as glycosaminoglycans (Mak and Mei, 2017). One of the most notable examples of BMs is in the intestine. Gut BM plays a crucial role as a semipermeable barrier and is highly responsive to both internal and exogenous stimuli. The dynamic interaction between structural components allow selective nutrient absorption while preventing transportation of harmful antigens and microorganisms (Vancamelbeke and Vermeire, 2017). Such complexity poses a challenge to the construction of a much-needed artificial bowel replacement substrate.

In this work, electrospun poly(ethylene terephthalate) (PET) was studied. PET is a type of nondegradable polyester, formed from condensation polymerisation of ester monomers which is a product from the reaction between ethylene glycol and terephthalic acid (Clark, 2004) **(Figure 1)**. It is widely used as plastics in food packaging and is FDA-approved (Custom-Pak, 2018). PET has desirable characteristics for tissue engineering such as biocompatibility, high tensile strength, chemical resistance and low cost (Subramaniam and Sethuraman, 2014). Existing literature shows promising results on PET as it already has successful biomedical applications such as in large vascular grafts and anterior cruciate ligament prosthesis (Subramaniam and Sethuraman, 2014).

**Figure 1.**
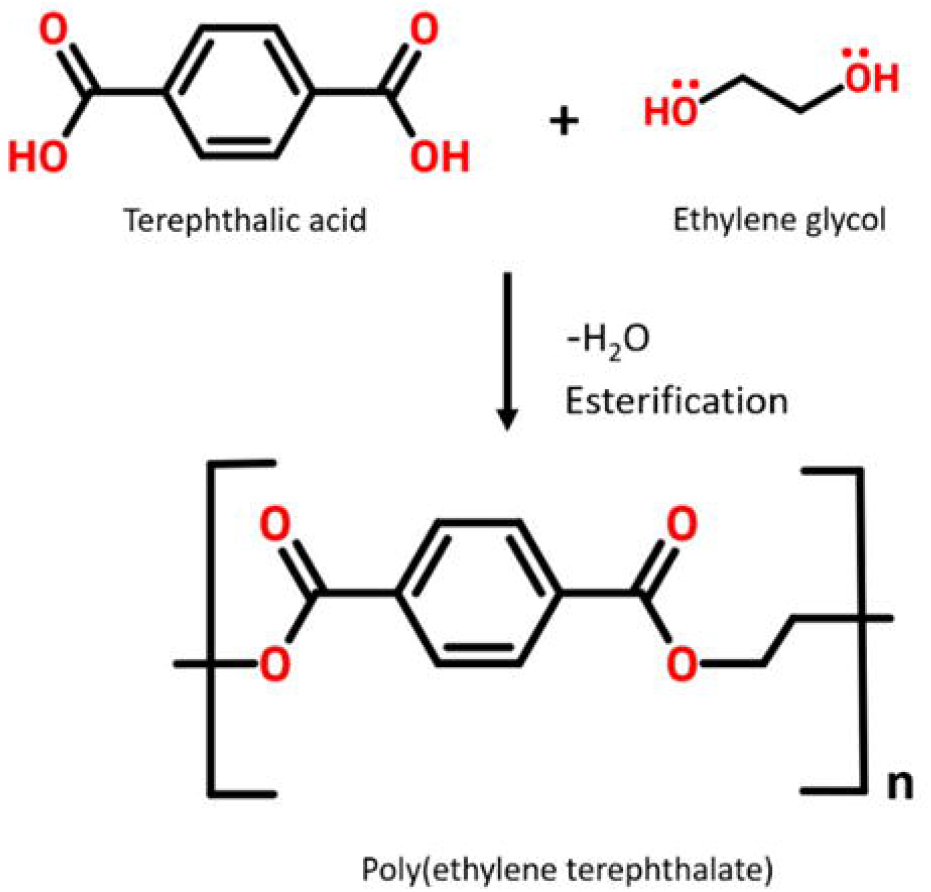
Schematic representation of the polymerisation reaction to form PET. Drawn using ChemWriter.

Electrospinning has gained significant popularity in the biomaterials field due to its simplicity and capability to extrude non-woven nanoscale fibres, allowing formation of polymeric membranes at a relatively low cost (Vass et al 2019). Electrospinning is initiated by application of electrostatic force to a droplet of liquid via high voltage supply. Once above a threshold voltage, the spherical droplet will elongate until it erupts into a conical shape referred to as Taylor cone (Agrahari et al., 2017). The electrostatic force overcomes the droplet surface tension, resulting in the extrusion of a charged liquid jet towards the grounded collector. The jet initially accelerates in a straight line but subsequently adapts a vigorous whipping motion which stretches and thins the resultant fibre. En route from the syringe tip to the collector, solvent in the polymer jet solution evaporates and fibres form a non-woven membrane on the collector surface (Xue et al., 2017). Fibre characteristics are influenced by multiple parameters that can be modified to suit its intended purpose, thereby rendering this technique as highly versatile and suitable for industrial-scale production (Kajdič et al., 2018). Spun fibres possess novel properties such as extremely high surface area-to-volume ratio, neat morphology and interconnected voids. These features propose electrospun membrane as the prime candidate for cell carrier scaffold due to its structural resemblance to natural extracellular matrix (ECM)(Jun et al., 2018) and its ability to promote native morphology and function in seeded fibroblast cell lines and small intestine organoids (Grikscheit et al 2004).

This article describes the optimisation process of an electrospun membrane and evaluation of its physical properties after various surface functionalisation treatments. The material discussed has the potential to be used as a permanent cell-carrier substrate that could contribute to the development of TESI, or for application in 3D models for small bowel research.

## Material and methods

### 1. Fabrication of PET fibre scaffolds

All experiments were undertaken in a custom-built grounded fume-hood using a custom-built electrospinning rig **(Figure 2)**. Pellets of PET (Polysciences Inc, USA) were dissolved in 1,1,1,3,3,3-hexafluoro-2-propanol (HFIP; Apollo Scientific, UK) at concentrations of 25%, 20%, 15% and 10% (w/v). After stirring for 24 hours at room temperature to ensure homogeneity, the polymer solution was introduced into a 10 ml syringe (BD Plastipak) and mounted onto a mechanical syringe pump to control the solution flow rate. A high-voltage power supply was used to apply a range of potential to the syringe. The electrospun fibre was deposited onto a grounded collector placed at various distances. All spinning parameters were summarised in **TABLES 1 and 2**.

**Table 1:** Summary of the range of PET/HFIP concentration and tip-to-collector distance used Tables

| Parameter | Range |  |  |  |
| --- | --- | --- | --- | --- |
| Concentration of PET/HFIP (%) | 25 | 20 | 15 | 10 |
| Tip-to-collector distance | 30, 25, 20, 15, 10 | 30, 25, 20, 15, 10 | 30, 25, 20, 15, 10 | 30, 25, 20, 15, 10 |

**Table 2:**
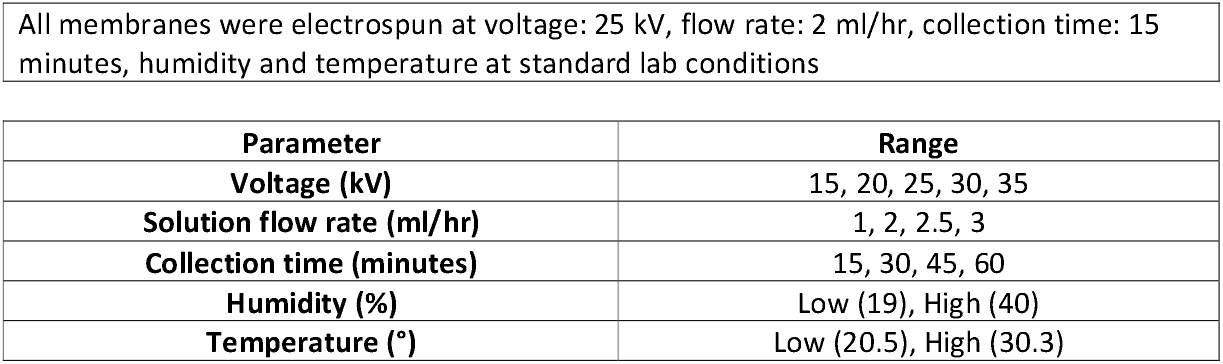
Summary of the range of operation parameters and environmental factors

| Parameter | Range |
| --- | --- |
| Voltage (kV) | 15, 20, 25, 30, 35 |
| Solution flow rate (ml/hr) | 1, 2, 2.5, 3 |
| Collection time (minutes) | 15, 30, 45, 60 |
| Humidity (%) | Low (19), High (40) |
| Temperature (°) | Low (20.5), High (30.3) |

**Figure 2.**
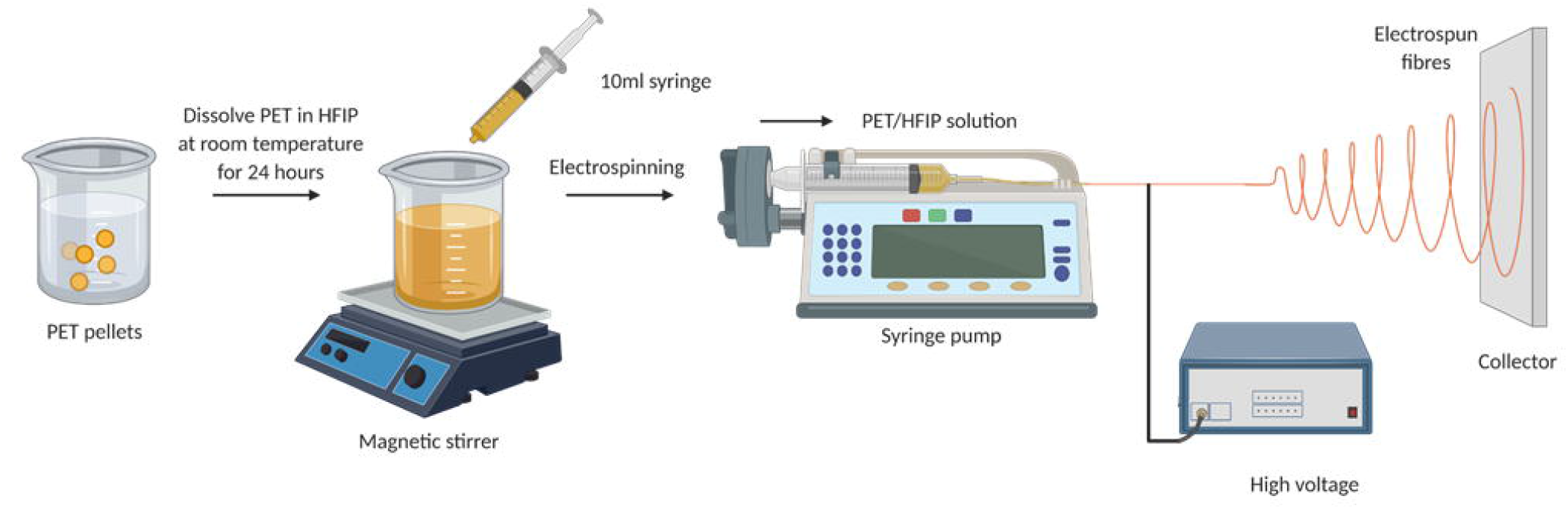
Schematic diagram of electrospinning setup. Drawn using BioRender.

Three membranes were electrospun for each parameter (n=3) except for 10% (w/v) at tip-to-collector distance of 10 cm where two membranes were fabricated (n=2).

### 2. Preparation of fibres for SEM

The surface morphology of the electrospun fibrous membrane was examined by scanning electron microscopy (SEM). Samples were attached to SEM stubs using carbon adhesive discs and then coated with 10 nm of platinum using a Q150T (QuorumTechnologies, Laughton, UK). Secondary electron SEM imaging was performed in a Quanta 250 FEG (FEI, Hillsboro, Oregon, USA) at low vacuum (0.53 Torr) with an acceleration voltage of 5 kV. Membranes were scanned at 5,000x and 10,000x magnifications. Images were analysed using ImageJ digital analysis software to measure the diameters of the individual fibres. A total of 50 measurements were taken at random and average fibre diameter calculated. Three membranes were analysed for each parameter (n=150) A fibre diameter distribution histogram was plotted using GraphPad Prism 8.3.0.

### 3. Mechanical testing of electrospun membranes

#### Quantitative analysis: tensile testing

Electrospun membranes were measured for tensile strength using a uniaxial tension/compression tester (Univert, CellScale) equipped with 1N load cell (Futek, California, USA) at a displacement rate of 2mm per minute. Membranes were manually cut into dog-bone shape*s* **(Figure 3.i)** to ensure the concentration of stress is in the middle of the sample when subjected to tensile force. The strips were subsequently mounted onto paper frames and the shoulder section taped down at each end. The paper frames were cut before tensile force applied. Membranes were tested until failure or for a maximum of 10 minutes, whichever came first. A digital calliper (Duratool) was used to measure the cross-sectional area of the fibre; these were used to calculate stress and strain. The gradient of the elastic region in the stress v strain curve **(Figure 3.ii)** equates to the tensile strength of the membrane.

**Figure 3.**
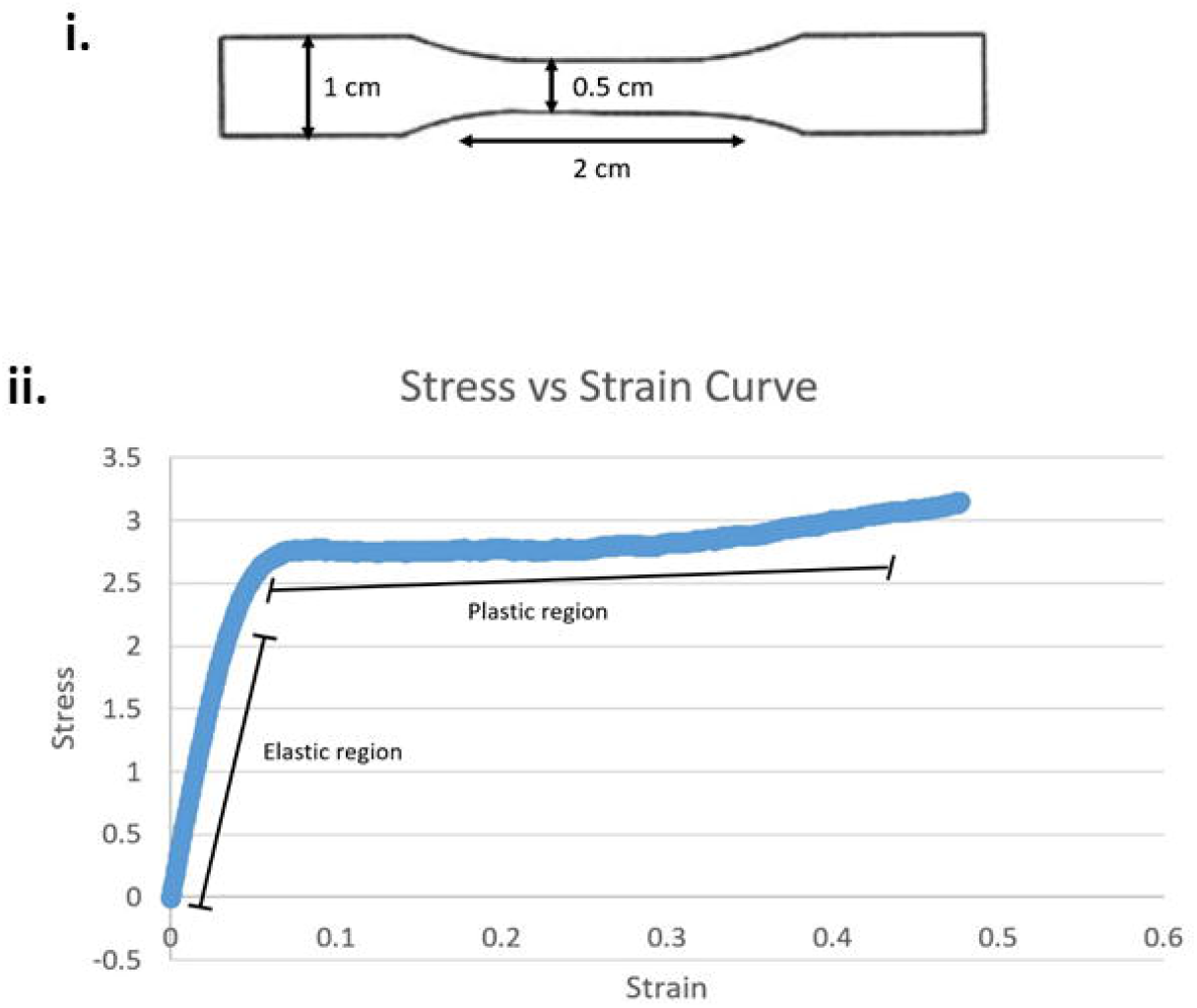
**(i)** Schematic diagram of dogbone sample dimensions **(ii)** Labelled stress vs strain curve showing elastic and plastic region

#### Qualitative analysis: manual handling

Membranes were folded, rolled and twisted to determine practicality of manual handling and observations noted.

### 4. Thickness and porosity

Membranes were cut into approximately 1 cm^2^ squares, thickness measured using a digital calliper (Duratool), and then weighed. Both measurements were performed three times on each membrane (n=3) and the porosity calculated using the following equation (Haneef and Downes, 2015):

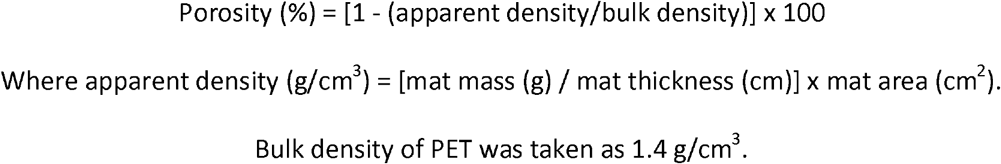

### 5. Contact angle measurements before and after surface treatment

Water contact angle (WCA) was measured using a drop shape analyser (DSA100, Kr*üss*, Hamburg, Germany). Three surfaces were analysed, including: tissue culture plastic as positive control, planar polymer surface and electrospun membrane. Planar polymer surfaces were fabricated by casting the polymer solution in a glass culture dish and left to evaporate overnight under fumehood flow to form a sheet. WCA was measured using a sessile drop method. A drop of 5µl distilled water was placed on the surface. Young’s contact angle was subsequently measured from an image of the sessile drop using *Kr*üss Drop Shape Analysis software. The angle was obtained at the three-phase boundary line between solid (membrane), liquid (water drop) and gas (air). All the WCA data were an average of three measurements at different locations on the surface (n=3).

### 6. Scaffold surface treatments

Multiple surface treatment modalities have been used in this study for biofunctionalization, particularly chemical and biological modifications including: ultraviolet (UV) irradiation, wet chemical methods and physical adsorption **(Supplementary Figure 1)**.

#### 6.1 UV treatment

The scaffold was positioned at a distance of 30 mm relative to the UV source (UVP CL-1000 Ultraviolet Crosslinker) and irradiated at the wave of 254 nm and energy of 15 mJ cm^-2^ for 15 or 30 minutes. Only one side of the membrane was treated; this was marked on the membrane to enable further characterisation of the treated side.

#### 6.2 Wet chemical and protein treatment

Two incubation time points (15 and 30 minutes) and two concentrations of sodium hydroxide (NaOH) solution (0.1 M and 1 M; Sigma) were examined. Electrospun PET membranes were fully submerged in NaOH solution at room temperature for the desired treatment time. The PET membranes were then washed three times in deionised water and left to dry on a non-stick sheet in laboratory conditions.

The same method was repeated for hydrochloric acid (HCl) solution (0.1 M and 1 M; Sigma), and fetal calf serum (FCS) diluted in PBS at 1 in 10 concentration and neat (LabTech).

### 7. SEM of membranes after surface treatment

10 × 10 mm PET membrane samples were attached to SEM stubs using carbon adhesive discs and then coated with chromium to reduce charging. SEM imaging was performed in a Hitachi S4800 high resolution SEM with an acceleration voltage of 2 kV. Membranes were scanned at 2,500x, 10,000x and 50,000x magnifications.

### 8. Mechanical testing of electrospun membranes after surface treatment

Identical setup to the tensile testing of membranes before surface treatment. Membranes were subjected to 10 cycles of 5 seconds of 0.3 mm displacement and 5 seconds of recovery, followed by stretch at a displacement rate of 3 mm per minute for a maximum of 10 minutes or until failure, which ever came first. A digital calliper (Duratool) was used to measure the cross-sectional area of the fibre; these were used to calculate stress and strain. The gradient of the elastic region in the stress versus strain curve equates to the Young’s modulus of the membrane. Three membranes were tested for each treatment condition (n=3)

### 9. Fourier Transform Infrared Spectroscopy (FTIR)

PET membranes were analysed using the diamond detector in the attenuated total reflection Fourier transform infrared spectroscopy (ATR-FTIR) (Bruker Vertex V70 FT-IR Spectrometer). The ATR-FTIR spectra were recorded between 400 and 4000 cm^-1^ and the ‘Peak Picking’ tool was used to identify peaks.

### 10. Permeability assay

A permeability assay measures the fluorescence intensity of fluorescein isothiocyanate (FITC) permeating through the PET membranes (Yeste et al. 2018). Untreated membranes and membrane treated with 1M NaOH for 30 minutes was mounted on cell-crowns and submerged in 800 µl of deionised water in a 24 well-plate. Subsequently 100 µl of 1:10000 (0.0185 mmol) FITC was pipetted onto the membrane, and the membranes were covered with foil and left for 10, 30 and 60 minutes. After the desired time point, three samples of the permeated FITC solution from each treatment group were pipetted into a 96 well-plate and analysed using the BMG FLUOstar OPTIMA Microplate Reader. Gain adjustment were performed to the fluorescence reading and converted into FITC concentration.

### 11. Statistical analysis

All statistical analysis was performed using GraphPad Prism 8.4.1 for Windows. Datasets were checked for normality using the Shapiro-Wilk test and statistical test performed accordingly. Data is presented as mean ± standard deviation. Statistical significance was taken at two-sided p-values of < 0.05.

## Results and Discussion

### Fibre morphology and fibre diameter

#### Qualitative analysis

Increase in fibre homogeneity with increase in polymer concentration was observed. Micrographs revealed the structural development from aligned arrays of smooth, continuous electrospun fibres at 25% (w/v) to beads on a string morphology at 15-20% (w/v) and distinct beading at 10%(w/v) **(Figure 4.i.)** As polymer concentration increased, the diameter of beads and distance between them increased. In addition, the beads transformed from spherical to ellipse-like shape **(Figure 4.i.B and Figure 4.i.D)**. These findings are consistent with existing literature and can be explained by polymer concentration, solution viscoelasticity and surface tension. Polymer concentration is directly proportional to solution viscoelasticity while surface tension depends on the nature of the polymer and solvent used (Fong et al., 1999). Due to surface tension, liquids display Plateau-Rayleigh instability to minimise their surface area; shrinking from streamline jets to sphere droplets (Haefner et al., 2015). Yu et al postulated that if viscoelastic forces of the polymer overcome surface tension then smooth fibres will be formed (Yu et al., 2006). Since the polymer and solvent used in this study and in turn the surface tension was kept constant, the degree of beading could be attributed to polymer concentration.

**Figure 4.**
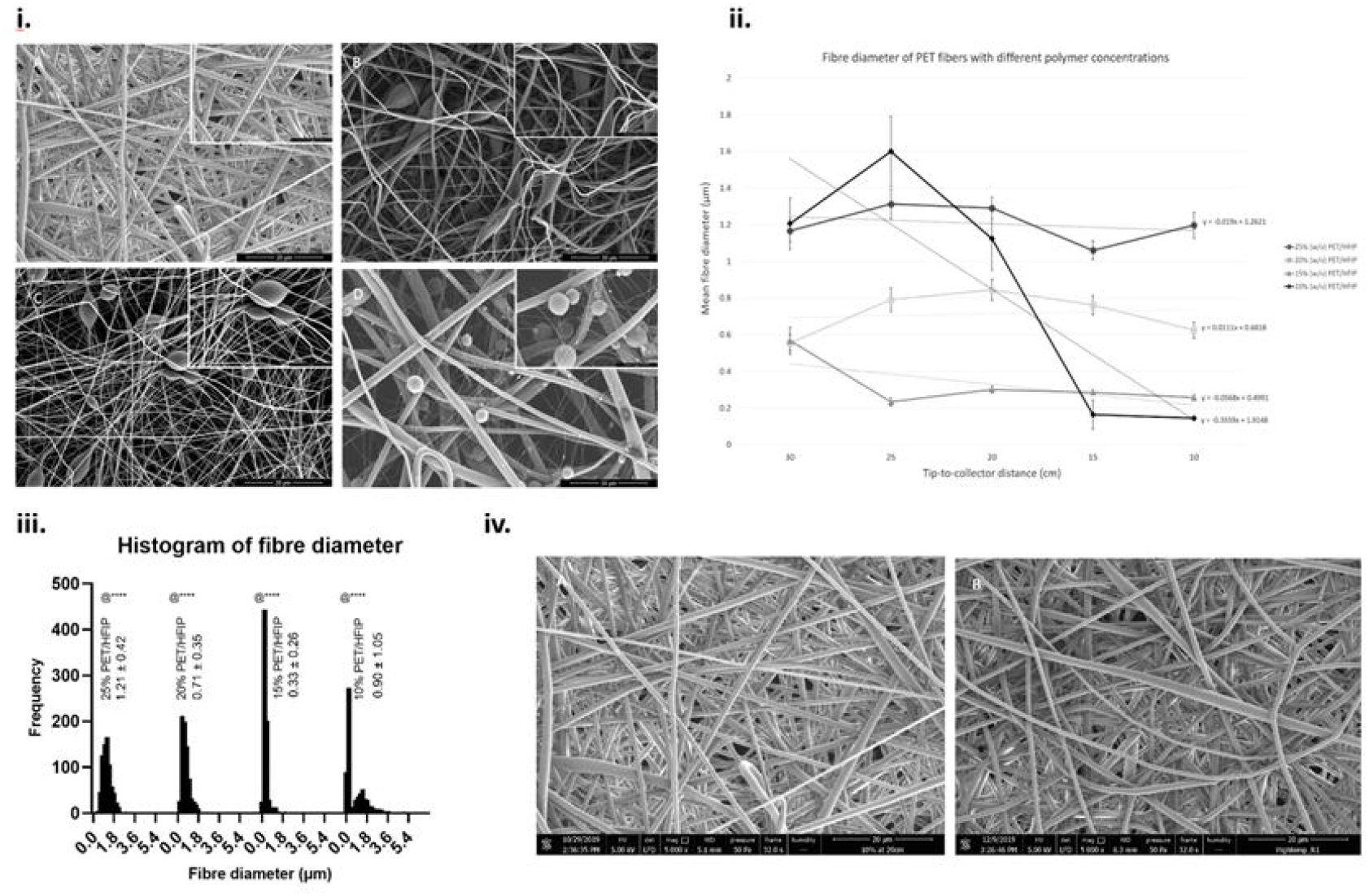
**(i)** SEM micrographs show polymer solution concentration affected fibre morphology. **(A)** 25% (w/v) PET/HFIP, **(B)** 20% (w/v) PET/HFIP, **(C)** 15% (w/v) PET/HFIP, **(D)** 10% (w/v) PET/HFIP; all fibres spun at tip-to-collector distance of 25 cm. Large image magnification: 5000x, small image magnification: 10 000x. Scale bar 20 µm. **(ii)** Scatter plot show polymer concentration had positive relationship with fibre diameter. All data presented as mean and CI. **(iii)** Frequency distribution graph of electrospun PET fibre diameter at different polymer concentrations. All data presented as mean ± SD. @ vs. all groups, ****=p<0.0001. n = 750 for concentrations of 25%, 20% and 15%, n = 700 for concentration of 10% **(iv)** SEM micrographs showing that temperature and humidity did not affect fibre morphology. **(A)** Low temperature and high humidity, **(B)** High temperature and low humidity. Magnification: 5000x. Scale bar 20 µm.

Qualitative analysis of the SEM images showed that the fibre morphology was not affected by some operation parameters **(Supplementary Figure 2)**. Experimental observations illustrated that membranes collected at 30 cm was significantly thinner and delicate compared to those collected at 10 cm; membranes collected at 10 cm were difficult to handle as it had the tendency to stick onto itself. Further, the proportions of the PET membrane sheet formed on the collector surface after the electrospinning process was also different. The size of the membrane sheet considerably decreased when the tip-to-collector distance decreased and became more circular in shape; corresponding with findings from Hekmati et al (Hekmati et al., 2013).

#### Quantitative analysis

In general, fibre diameter was found to increase with polymer concentration **(Figure 4.ii)**. This is due to higher output of polymer extruded at any one time with higher concentrations. However, fibres obtained from 10% (w/v) membranes had a larger average diameter than 15% and 20% (w/v) fibres as well as bigger standard deviation. As aforementioned, low solution concentration and therefore low viscoelasticity led to unstable electrospinning, resulting in high variation of electrospun fibres. In addition, 10% (w/v) fibre diameters showed bimodal distribution **(Figure 4.iii)** therefore the mean may not be representative of the whole population. No significant relationship between tip-to-collector distance and fibre diameter could be concluded. This finding is in contrast with previous literature which reported a positive correlation and stated the increase in the needle-tip-to-collector distance allows the polymer solution jet to bend and whip more extensively, thus decreasing fibre diameter (Mazoochi et al., 2012).

Solution flow rate of 1 ml/hr significantly reduced fibre diameter. ANOVA analysis of the flow rate parameter showed significant difference among means (F = 8.232, df = 599, p<0.0001) and Tukey post hoc analysis illustrated that fibres spun at 1 ml/hr had significantly smaller diameter than those spun at other flow rates. However, other flow rates did not display significant differences **(TABLE 3)**. Flow rate directly influenced the amount of solution present at the syringe tip at any one time; therefore, low flow rate means less extrusion of polymer solution, resulting in small diameter.

**Table 3:** There were no significant correlation between average fibre diameter in each parameter. All data presented as mean ± SD.

| Operation parameter | Range |  |  |  |  |
| --- | --- | --- | --- | --- | --- |
| Voltage (kV) | 35 | 30 | 25 | 20 | 15 |
| Fibre diameter ($\mu\text{m}$) | $0.947 \pm 0.347$ | $1.32 \pm 0.363$ | $1.31 \pm 0.501$ | $0.884 \pm 0.513$ | $1.31 \pm 0.498$ |
| Flow rate (ml/hr) | 3 | 2.5 | 2 | 1 |  |
| Fibre diameter ($\mu\text{m}$) | $1.27 \pm 0.471$ | $1.35 \pm 0.336$ | $1.31 \pm 0.501$ | $1.12 \pm 0.319$ | |
| Collection time (mins) | 60 | 45 | 30 | 15 |  |
| Fibre diameter ($\mu\text{m}$) | $1.22 \pm 0.376$ | $1.24 \pm 0.343$ | $1.33 \pm 0.412$ | $1.31 \pm 0.501$ | |

Interestingly, spinning voltage, solution flow rate and collection time did not affect fibre morphology or fibre diameter. This is inconsistent with existing literature that states spinning voltage directly influences the stretching force of the polymer jet, which in turn regulates thinning of the jet (Lee et al., 2004, Buchko et al., 1999). Hence, an increase in voltage leads to thinner fibres. However, Zhao et al. demonstrated that beyond a threshold value, high voltage may result in thicker fibres due to reduction in flight time (Zhao et al., 2004). Multiple groups have reported an increase in fibre diameter with increase in solution flow rate since there is an increase in volume of polymer being syringed out at any one time (Zargham et al., 2012, Milleret et al., 2011). Further, an increase in flow rate may imply emergence of secondary jets; during the electrospinning process, the initial jet can solidify in certain places on the nozzle tip, leading to eruption of multiple secondary jets from unsolidified areas. These secondary jets each contain smaller volumes of polymer solution than the initial jet, resulting in smaller fibres (De Schoenmaker et al., 2012). This phenomenon was observed during electrospinning in our study. Additionally, increase in collection time may increase the possibility of solidification of the initial jet, leading to a decrease in the average fibre diameter.

Effect of environmental parameters of temperature and humidity was tested on 25% (w/v) PET membranes because it showed defect-free fibre morphology at standard laboratory conditions. Temperature and humidity did not affect fibre morphology and fibres were defect-free **(Figure 4.iv)**. However, larger fibre diameters were produced when the temperature was high and humidity was low. Unpaired t-test of fibre diameters showed significant difference between means of 0.211 ± 0.0578 (F = 1.001, df = 298, p-value = 0.0003). This indicated fibres spun at high temperature and low humidity have significantly larger average diameter. Currently, there is contradicting evidence in literature on the effect of temperature and humidity on fibre diameter. Wang et al. (Wang et al., 2007) reported that temperature elevation was an effective method to produce ultrathin fibres whilst works from Guangzhi et al. (Guangzhi et al., 2017) suggested that excessively high temperature can lead to large fibres. Higher humidity resulted in larger polystyrene fibre diameter (Kim et al., 2004), but Htike et al. (Htike et al., 2012) demonstrated fibres were thinner instead. These inconsistent findings may be attributed to multiple competing factors on the electrospinning process. An increase in temperature will decrease solution viscosity and surface tension (Guangzhi et al., 2017), resulting in smaller fibres. Low humidity can lead to reduction in precipitation of the polymer when water condenses on the surface of the polymer jet, resulting in thinner fibres (Ogulata and İçoğlu, 2013). Both high temperature and low humidity directly accelerate solvent evaporation from fluid jets. Premature evaporation can lead to reduced stretching of the polymer fluid jet, thus fibres with large diameters were produced (Guangzhi et al., 2017). Golin et al. postulated that during the electrospinning process, fibres take on moisture from the surrounding atmosphere; decrease in humidity leaves less water vapour present in the atmosphere, leading to less water absorption, hence fibres solidify more rapidly allowing less time for stretching (Golin, 2014). Our study included only two combinations of temperature and humidity variant, therefore, further experiment is needed to ascertain the effects of environmental factors on PET fibre diameter.

For the following experiments both 20% and 25% (w/v) PET membranes were investigated to determine difference between defect-containing and defect-free membranes.

### Mechanical testing and handling

20% PET membranes collected at the tip-to-collector distance of 30 cm exhibited lowest tensile strength **(Figure 5.i.)**. There were no significant differences among other membranes. However, existing literature on poly(ε-caprolactone) materials demonstrated a positive correlation between collector distance and tensile strength (Asvar et al., 2017); increasing distance results in longer travel time to the collector, therefore the solution is able to undergo extended whipping, theoretically resulting in thinner and greater number of fibres per area. This results in fibres being more tightly packed leading to increased mechanical strength. This contradiction indicates that effects of electrospinning parameters on an output might be dependent on the polymer composition. Qualitative analysis of membranes during tensile testing showed both 20% and 25% (w/v) membranes formed at tip-to-collector distance of 25 cm stretched most uniformly and did not reach point of failure after 10 minutes **(Figure 5.ii.)**. Therefore, membranes spun with polymer concentration of 20 and 25% and collected at tip-to-collector distance of 25 cm were taken forward for further characterisation.

**Figure 5.**
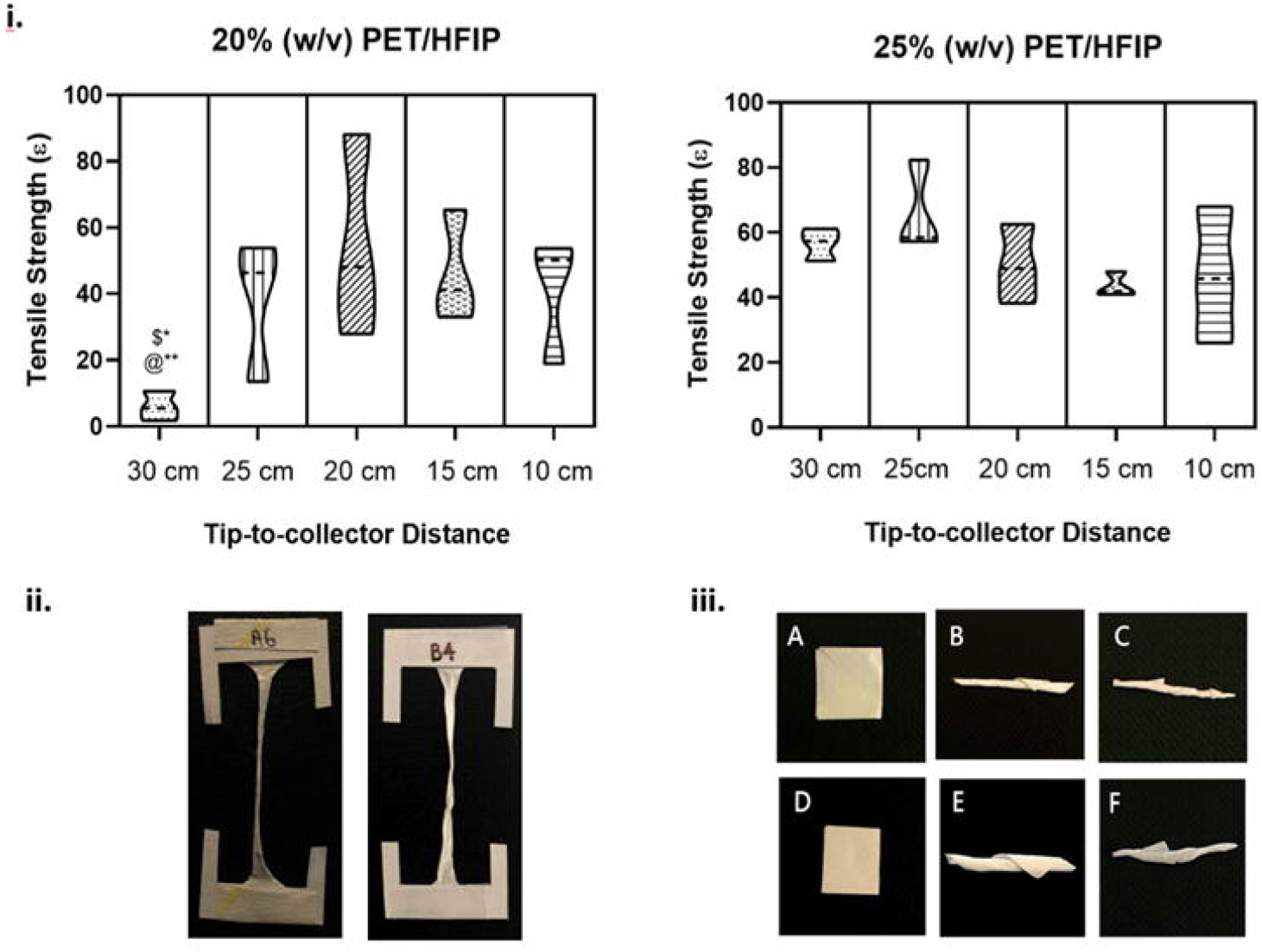
(i) Violin plot show 20% (w/v) membranes collected at 30 cm had the lowest tensile strength. Violin plot indicates upper and lower quartiles and middle dotted line represents the median. ANOVA: F=2.634, df = 29, p-value< 0.05. Tukey post hoc: $ vs. 25% at 30 cm, @ vs. 25% at 25 cm. * = p<0.05, ** = 0.001≤p ≤0.01 **(ii)** Photographs show intact PET membranes stretched uniformly after tensile testing. **(A)** 25% (w/v) collected at 25 cm, **(B)** 20% (w/v) collected at 25 cm **(iii)** Photographs of PET membranes undergoing **(A),(D)** folding, **(B),(E)** rolling and **(C),(F)** twisting without being damaged. **(A)-(C)** are 20% PET/HFIP fibres, **(D)-(F)** are 25% PET/HFIP fibres

Both 20 and 25% (w/v) PET membranes could withstand folding, rolling and twisting equally well without being damaged **(Figure 5.iii)**.

### Thickness and porosity

25% (w/v) electrospun membranes were thicker and more porous than 20% (w/v) membranes **(TABLE 4)**. Higher polymer concentration led to thicker membrane due to production of thicker fibre and looser packing, resulting in larger voids between the fibres. High membrane porosity could also be explained by the same theory.

**Table 4:** 25% (w/v) membranes are thicker and more porous than 20% (w/v) membranes. Both electrospun at tip-to-collector distance of 25 cm. All data presented as mean ± SD.

| Fibre parameters | Thickness ($\mu\text{m}$) | Porosity (%) |
| --- | --- | --- |
| 20% PET/HFIP at 25 cm | $50.0 \pm 8.16$ | $89.3 \pm 2.9$ |
| 25% PET/HFIP at 25 cm | $76.7 \pm 9.43$ | $92.6 \pm 1.9$ |

### Wettability

Untreated electrospun PET membranes exhibited poorer wettability versus untreated planar PET and tissue culture plastic (TCP) (ANOVA: F=86.9, df=14, p<0.0001), in both defect-free (25% (w/v)) and defect-containing (20% (w/v)) membranes **(Figure 6)**. Planar PET surfaces represent the inherent surface chemistry of PET as it had not been influenced by the electrospinning process. WCA of untreated planar PET was similar to that of TCP; the positive control widely used for cell culture. Our findings suggested that innate surface chemistry of PET was indeed hydrophilic, which could be attributed to oxygen molecules present in the PET polymer. Therefore, poor wettability was due to surface topography. Electrospun membranes are an open network of voids that exhibit rough surfaces leading to formation of air pockets between the drop and the membrane, a phenomenon referred to as Cassie-Baxter wetting state (SUSANNA LAURÉN). However, the inherent hydrophilic chemistry of the polymer would remain, and given the high surface area due to the membrane’s morphology would expose more of the chemical structure on a molecular level. Biomaterial wettability has implications in cell attachment and interaction towards the surface of any given material. Webb et al. reported that cell attachment was significantly greater on hydrophilic surfaces relative to hydrophobic surfaces (Webb et al., 2000). Electrospun membranes can be modified to become more hydrophilic using treatments including, but not limited to, UV/ozone, weak acid, weak alkali and protein attachment e.g. bovine serum albumin.

**Figure 6.**
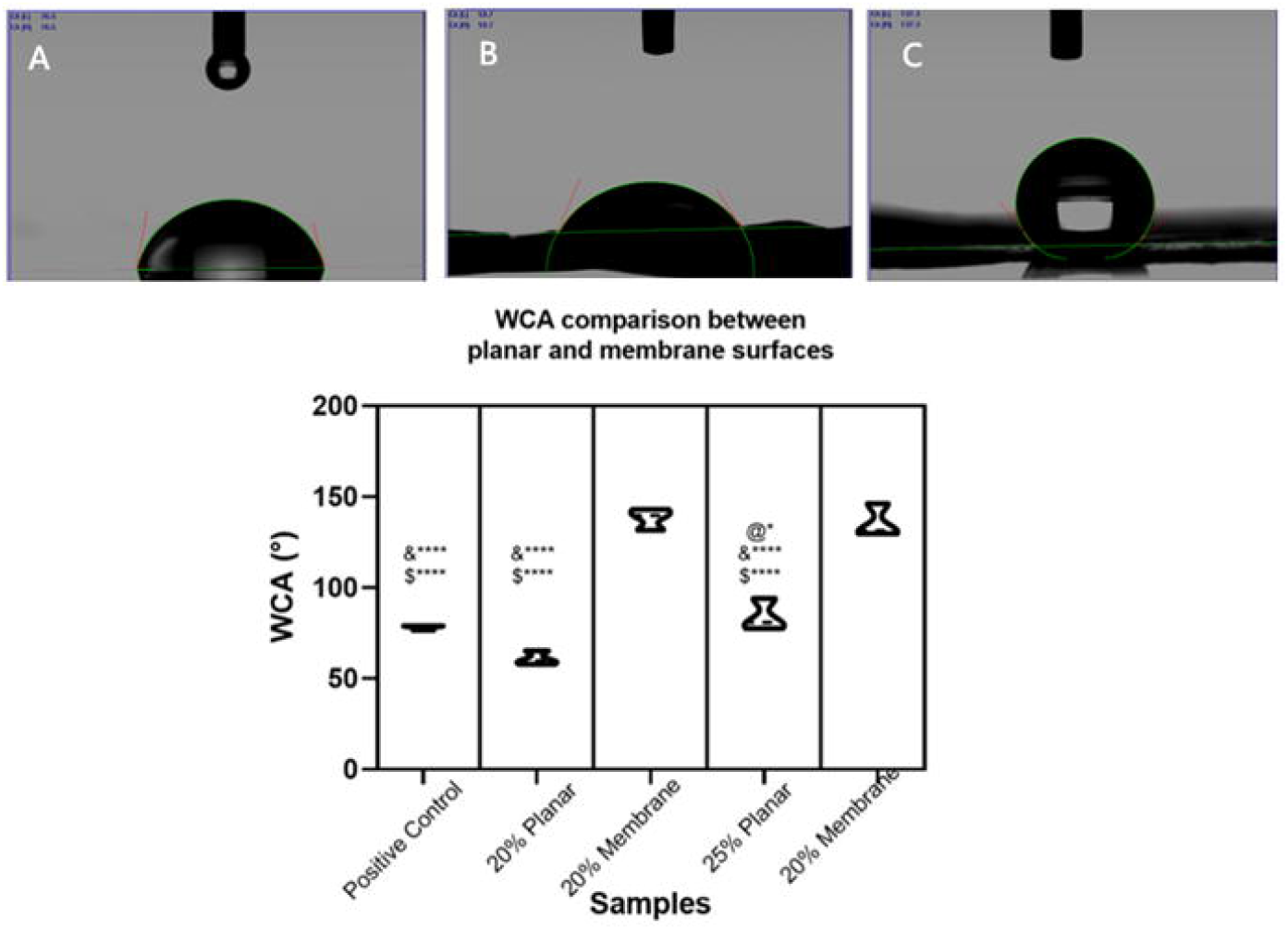
PET membranes were more hydrophobic than planar surfaces and tissue culture plastic. Images comparing WCA between positive control, planar surfaces and fibres made from 20% and 25% (w/v) PET/ HFIP concentrations. **(A)** WCA on treated polystyrene, **(B)** WCA on planar surface **(C)** WCA on membrane surface. Violin plot indicates upper and lower quartiles and middle dotted line represents the median. & vs. 20% fibre, $ vs. 25% fibre, @ vs. 20% planar. * = p<0.05, **** = p<0.0001

We undertook further surface modification tests on defect-free membranes only (25% (w/v)) to ensure that the only variables were: surface treatment type (UV, acid, alkali, or FCS) and treatment time (15 minutes and 30 minutes).

### Wettability & ATR-FTIR after surface treatments

Significant improvement in hydrophilicity was observed in membranes treated with NaOH 0.1 M 30 minutes, 1 M NaOH and diluted FCS irrespective of treatment times **(Figure 7)**. Lowest WCA (0° ± 0°) was denoted for membranes submerged in NaOH 1 M 30 minutes, as the water drop did not persist long enough on the treated surface to measure. Contact angle of NaOH-treated samples, decreased with increased concentration and submersion time. Mean WCA of membranes treated with diluted FCS also reduced with increased treatment time. No significant change was observed with increased FCS concentration, nor with UV or HCl treated membranes.

**Figure 7.**
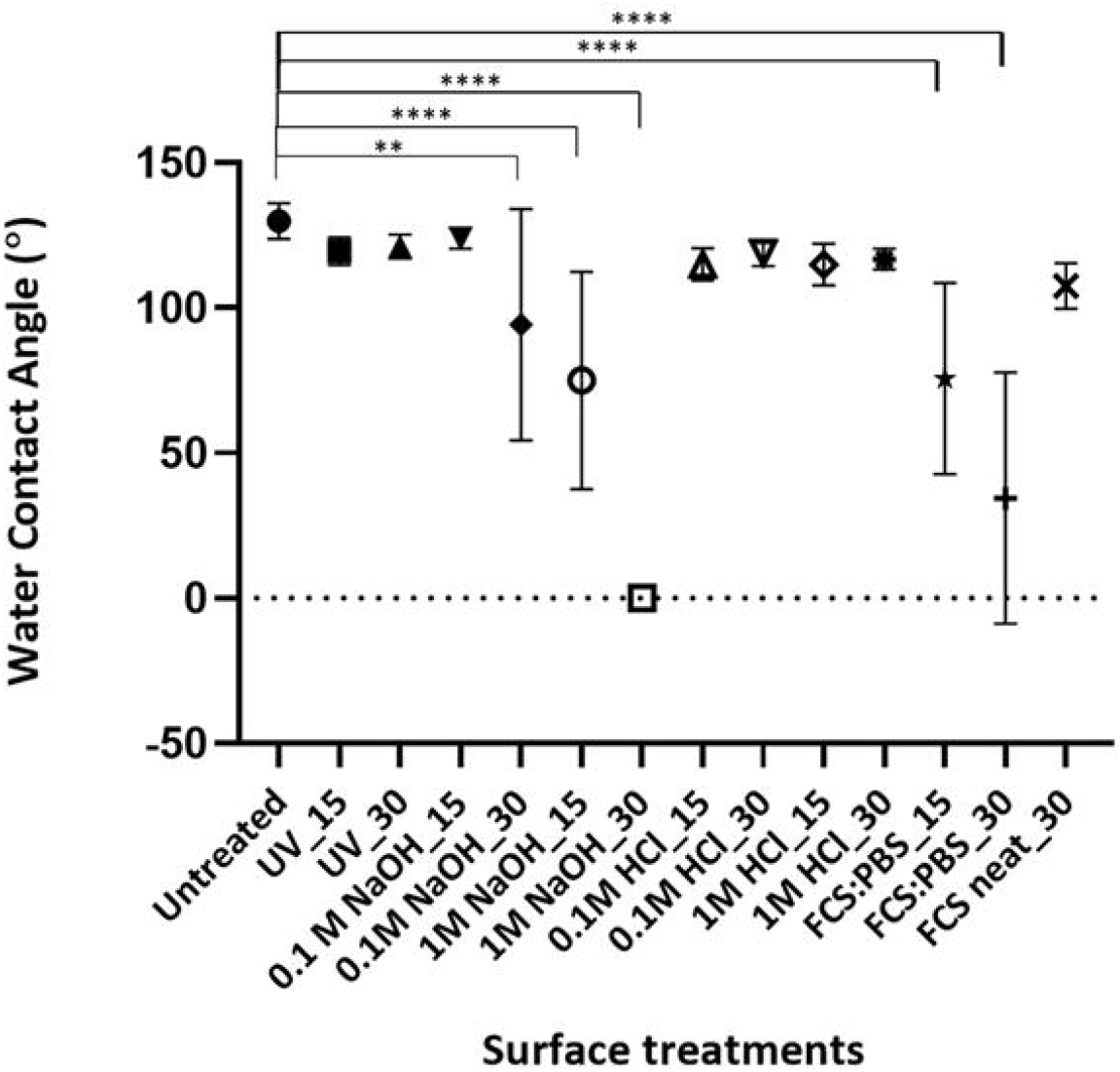
Membranes treated with 0.1 M NaOH for 30 minutes, 1 M NaOH and diluted FCS show significant reduction in WCA. WCA measurement of treated membranes to determine degree of hydrophilicity. Lowest WCA observed in 1 M NaOH for 30 minutes treatment group. Data are shown as mean ± standard deviation, n=9. Two-way ANOVA: F_(13, 104)_ = 29.6, df = 13, p-value <0.05. Dunnett’s multiple comparisons test: ** p=0.005, **** p<0.0001. Surface treatments written as ‘Treatment_Submersion time’.

Hydrophilicity is influenced by three main factors (Haneef and Downes 2015, Polini and Yang 2017). Firstly, membrane surface chemistry governs the interactive forces between polar functional groups in the polymer surface with the polar water molecules. Secondly, fibre topography and structural architecture effects air trapping between the membrane pores, which can alter the interface between droplet and membrane surface. Lastly, the size of the water droplet can affect hydrogen bonding and the degree of droplet spread. As the droplet size was kept constant at 5 µl, this factor will not be discussed. Multiple groups regard WCA values less than 90° to be hydrophilic (Purkait, Sinha et al. 2018), therefore all untreated PET membranes were hydrophobic (130° ± 6.17). Theoretically, the presence of ester groups in the polymer backbone should promote hydrophilicity but this may have been counteracted by the non-polar phenyl cyclic hydrocarbon ring and hydrocarbon backbone that repels the polar water molecule. In addition, surface roughness as a result of electrospun topography and random fibre orientation promotes air trapping between membrane voids, increasing WCA.

Figure 7. illustrated improvement of hydrophilicity in membranes treated with NaOH to be a concentration- and submersion-time-dependent process. Treating membranes with NaOH 1 M 30 minutes turned it into a superwetting material. NaOH solution performs random chemical scission on ester linkages, resulting in exposure of hydroxyl and carboxylate groups on the polymer surface. These functional groups are able to form hydrogen bonds with water molecules, and in turn improve hydrophilicity. ATR-FTIR exhibited the change in surface functional groups following treatment with NaOH and HCl; untreated membranes showed strong ester bonds as expected and was correctly identified as PET **(Figure 8a)**. Both wet chemical treatments **(Figure 8b-c)** exhibited a stronger hydroxyl group peak than the untreated membrane due to hydrogen bonds in the carboxylate group and secondary alcohols. Both spectra also showed more C–H stretching as a result of chemical scission.

The infrared spectra of NaOH-treated membrane **(Figure 8b)** correlated with reduction in WCA as it showed the presence of hydrophilic hydroxyl group. This is shown by an increased intensity of bands in the 3200 – 3550 cm^-1^ region and the 2960 cm^-1^ peak compared to the untreated membrane. Although exhibiting similar spectra, HCl-treated membranes did not demonstrate significant reduction in WCA, which correlated with a much lower detection intensity in the aforementioned peaks **(Figure 8c)**. This may be because hydrolysis of esters catalysed by acid is much slower at any given temperature than alkali-catalysed hydrolysis (Mashiur 2012). It may be that the small alteration in surface chemistry that was detected by ATR-FTIR for HCl treated membranes was not significant enough to translate to a change in physical properties.

**Figure 8.**
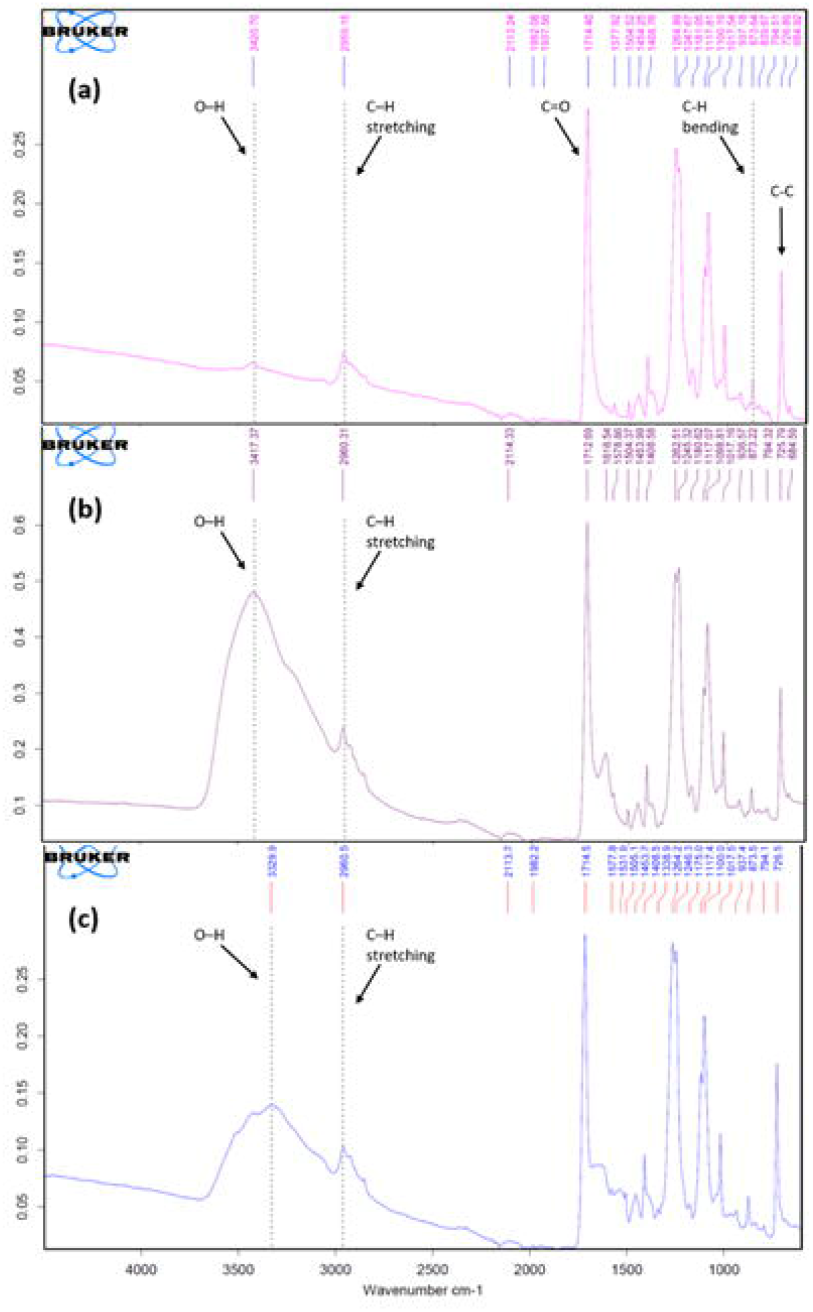
Membranes treated with HCl and NaOH showed more intense –OH bands and C– H stretching than untreated membranes. ATR-FTIR spectra was used to analyse the surface chemistry of PET membrane. (a-c) are representative spectra of membranes treated with wet chemical method: (a) untreated, (b) treated with HCl and (c) treated with NaOH.

Submerging membranes in diluted FCS solution promotes protein adsorption. The protein attaches via interactions such as weak Van der Waal forces, electrostatic forces, hydrogen bonds and hydrophobic dehydration (Tallawi M 2015). Hydrophobic dehydration happens between hydrophobic regions on the surface and hydrophobic domains on the protein structure. This interaction promotes protein conformational change and unfolding. Regardless of protein original structural arrangement, such unfolding increases the exposure of active sites for protein surface contacts (Felgueiras, Antunes et al. 2018). Membranes treated with neat FCS had higher WCA than the diluted FCS group and had no significant change compared to the untreated group. Protein adsorption is complex and depends on many factors. Firstly, many proteins exist in FCS and the Stokes-Einstein-Sutherland equation predicted that small proteins will diffuse faster than larger proteins. Secondly, solution concentration influences the extent of protein saturation on the material surface. If the surface is sub-saturated, then there is available space for proteins to unfold and favourable conformation may be achieved, whereas proteins on saturated surfaces must displace adsorbed neighbours, slowing down the unfolding process. At surface saturation (which may be the case in the neat FCS group), smaller proteins that arrived quickly has already formed a monolayer coverage. Larger proteins that reach the membrane subsequently, then forms multi-layer absorption on top of the smaller proteins. However, this form of adsorption is weaker and larger proteins can be washed off during the rinse with deionised water immediately after treatment. Therefore, membranes treated with neat FCS may have less attachment of larger proteins compared to membranes submerged in diluted FCS. As larger proteins contain more functional groups and can contribute to membrane hydrophilicity more significantly than smaller proteins, the reduction in WCA is therefore limited in the neat FCS group (Vogler 2011, Felgueiras, Antunes et al. 2018). ATR-FTIR for FCS treated membranes exhibited no change in surface chemistry (data not shown).

There was no significant reduction of WCA of membranes after UV irradiation. This is conflicting with existing evidence that shows drastic improvement in hydrophilicity (from 83.8° ± 1.0 to 40.8° ± 1.4) in PET films after treatment with UV excimer light at 15.8 mJ cm^-2^ for 2 - 90 seconds (Gotoh, Yasukawa et al. 2011). Reduction in WCA was attributed to the introduction of oxygen-containing groups onto the film surface after UV treatment, leading to an increase in the base parameter of the surface free energy. UV treatment also results in the formation of low-molecular-weight oxidised materials (LMWOM) on the polymer surface, arising from random scission of polymer chain to form hydrophilic oligomers (Gotoh, Yasukawa et al. 2011). However, these hydrophilic groups can undergo surface rearrangement and be lost to the atmosphere over time in a process called ‘hydrophobic recovery’, hence losing its hydrophilicity. In this work, WCA was measured 1-week post-treatment. This coincides with the time-point whereby hydrophobic recovery reaches stability in the literature, which may explain the lack of WCA reduction post UV irradiation. Gotoh et al. suggested rinsing the membrane with water immediately after treatment may help suppress the extent of hydrophobic recovery. ATR-FTIR for UV treated membranes exhibited no change in surface chemistry (data not shown).

Although there is evidence that hydrophilic scaffolds can improve cell adhesion, extreme hydrophilicity may not be desirable. Cell adhesion studies carried out using human osteosarcoma cell on polystyrene scaffold concluded that optimal adhesion was observed at WCA of 64°(Dowling, Miller et al. 2011). This correlates with a study on low density polyethylene surface, which demonstrated stronger adhesion force at WCA approximately 60 - 65° (Xu and Siedlecki 2007). Membranes with mean WCA closest to this range are those treated with NaOH 1 M 15 minutes (75.1° ± 37.4) and diluted FCS 15 minutes (75.6° ± 33.0).

### Fibre morphology and diameter after surface treatments

Figure 9. shows representative SEM images of fibre morphology in each treatment group and the corresponding histogram plot of fibre diameters. All surface treatments resulted in morphological change to varying degrees, except for FCS (data not shown). 1M NaOH-treated membranes appears to have the roughest morphology and those treated with HCl underwent the least change. Rough morphology in most treatment groups was limited to the surface, however fibres treated with 1 M NaOH, irrespective of submersion time, showed deeper degradation and salt crystal deposition. NaOH attacked a large proportion of the fibre surface, causing significant surface etching and extends inside the fibre, leading to formation of elongated cavities on the surface **(Figure 9(g))**.

There is evidence that cell adhesion and cell proliferation is improved on rough surfaces and thinner fibres (Anselme 2000, Xu, Yang et al. 2004, Wan, Wang et al. 2005) (Kurokawa, Endo et al. 2017), however treatment parameters must be balanced to minimise compromise to mechanical integrity. Furthermore, existing literature reported that rougher surfaces give higher WCA (Cui, Li et al. 2008). However, this contradicts with the results from our study as NaOH-treated membranes resulted in lowest WCA values. Membranes treated with NaOH 1 M 15 minutes and those treated with HCl 0.1 M 30 minutes both resulted in significant reduction in mean diameter compared to untreated membranes; 30.6% and 22.1% decrease, respectively. Previous work has shown that NaOH can lead to fibre thinning (Musale and Shukla 2016) and membrane weight reduction (Lee, Park et al. 2012) but there is lack of such evidence for HCl.

**Figure 9.**
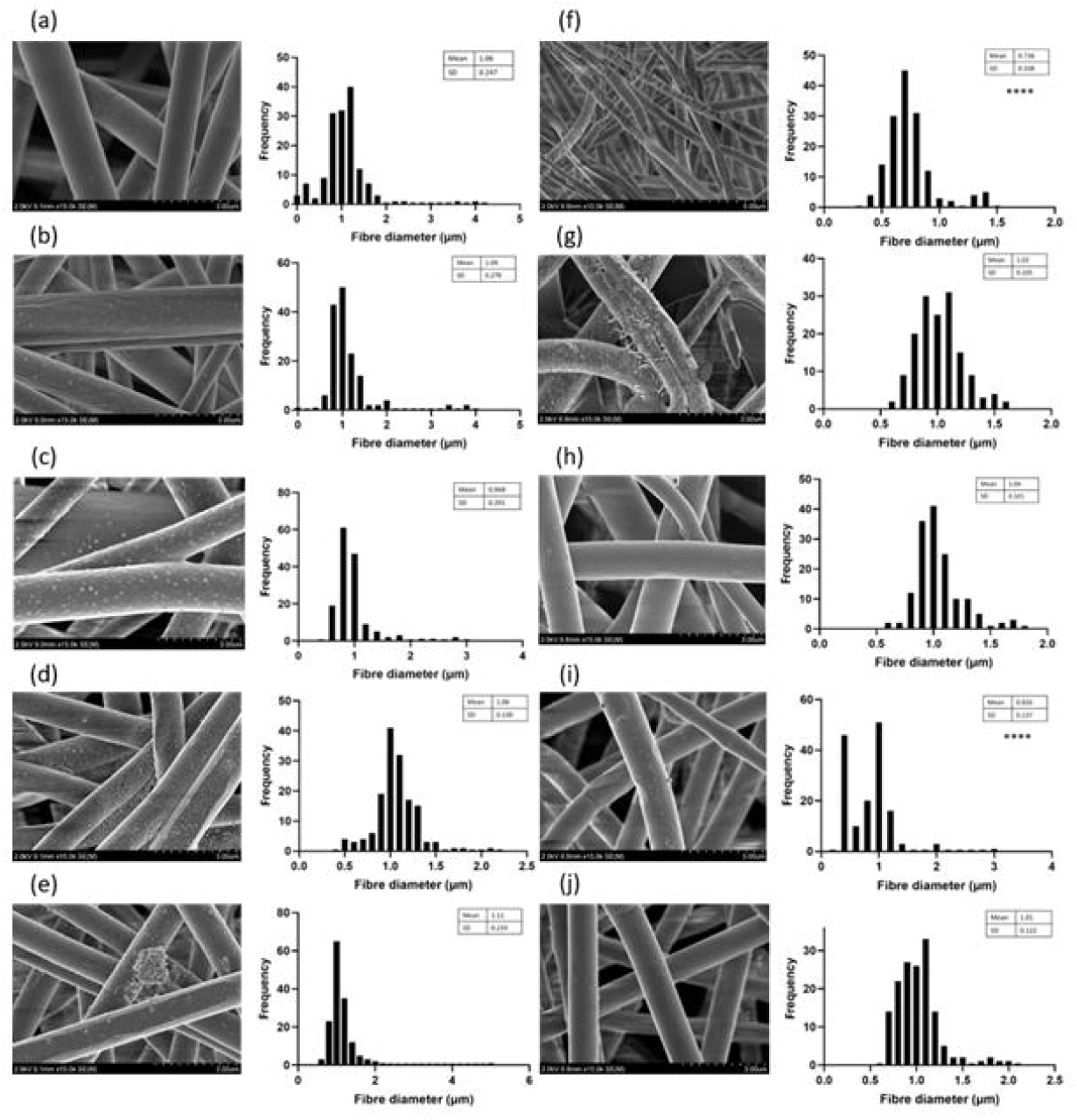
Every surface treatment condition led to fibre morphological change. Membranes treated with 1 M NaOH for 15 minutes and membranes treated with 0.1 M HCl for 30 minutes both had reduced fibre diameters. Representative SEM images demonstrating the effects of different surface treatments on membrane morphology (magnification x15,000, except (f) x10,000) and histogram plot of fibre diameter frequency distribution (n=150). (a) untreated, (b) UV for 15 minutes, (c) UV for 30 minutes, (d) 0.1 M NaOH for 15 minutes, (e) 0.1 M NaOH for 30 minutes, (f) 1 M NaOH for 15 minutes, (g) 1 M NaOH for 30 minutes, (h) 0.1 M HCl for 15 minutes, (i) 0.1 M HCl for 30 minutes and (j) 1 M HCl for 30 minutes. Kruskal-Wallis: p<0.0001. Dunnett’s multiple comparison: **** = p<0.0001

### Mechanical properties after surface treatments

The scatter plot shown in **Figure 10.i**. is a representative plot illustrating the relationship between force and displacement during 10 cycles of stretch and relaxation of membranes. There is no statistical difference between each cycle (p = 0.966) which means that the membrane did not lose its elastic property during repeated cyclic tensile test. This is true for membranes in every treatment group. It can be seen that the ‘stretch’ phase of the first cycle required more force to achieve the same displacement as subsequent cycles. The graph for most samples followed this trend, except membranes treated with 0.1 M HCl where there was no observable distinction between the initial cycle and the subsequent cycles.

**Figure 10.**
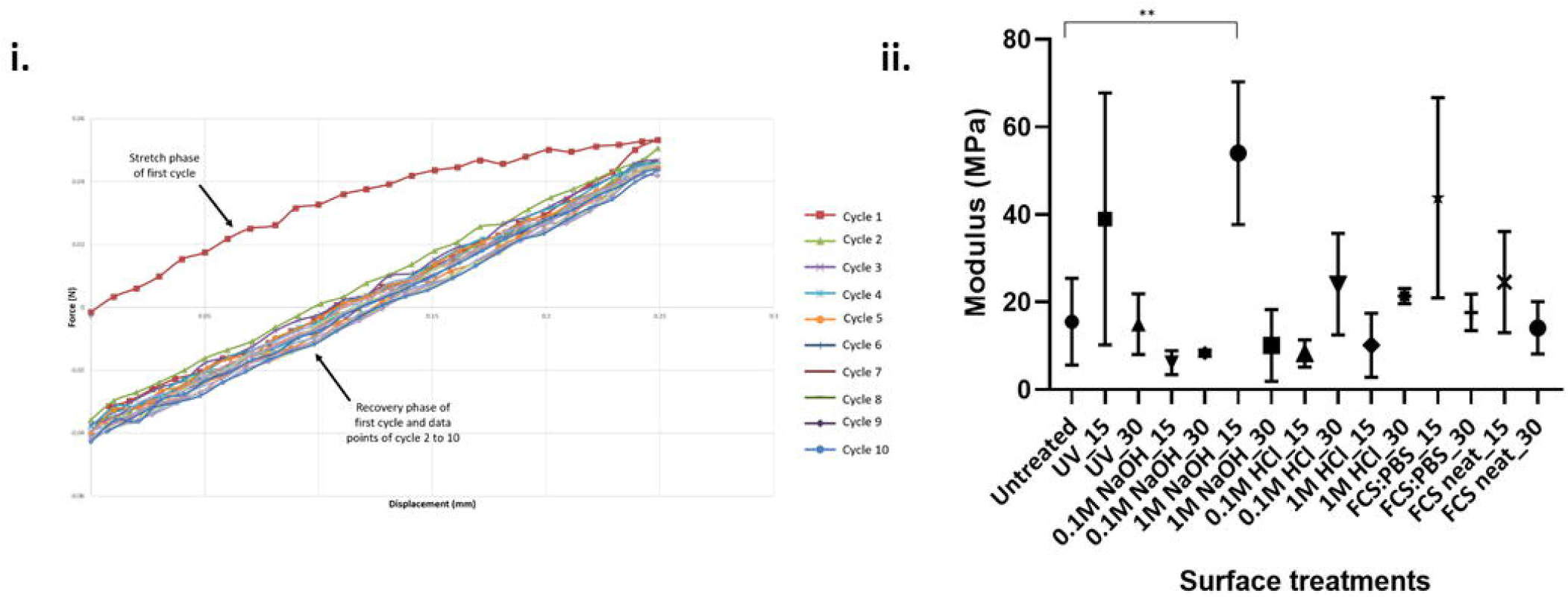
Membranes in every treatment group were able to retain its elastic property during repeated cyclic tensile test. **(i)** Membranes were subjected to 10 stretch-relaxation cycles during tensile testing to evaluate its elasticity. Representative force vs displacement scatter plot demonstrates similar trend across all cycles. Gradient of the stretch phase of each cycle was subjected to statistical analysis. One-way ANOVA: F_(9, 20)_ = 0.300, df = 9, p-value = 0.966. **(ii)** NaOH 1 M 15 minutes treatment group exhibited significantly higher modulus than untreated membranes.

On a molecular level, during the relaxed state the polymer chains are coiled and when stretched the chains are extended, acting like springs. As long as the uniaxial load applied is not sufficient to break the chemical bonds, the membrane will return to its original form. Although not statistically significant, the initial stretch phase can be seen to follow a different trend compared to subsequent cycles and require higher force to achieve the same displacement. This initial force might have been used to break interactions between polymer chains that kept the chains ‘coiled’, but insufficient to disrupt intermolecular bonds that will render the membrane plastic. Further, the stretch phase of the first cycle follows the Maxwell model of viscoelasticity (Schiavi and Prato 2016); when the material is subjected to constant strain, the stresses gradually relax. This is shown by the shape of the graph whereby the force tapers off as it reaches maximum displacement. Meanwhile, the graph of subsequent cycles follows a near-perfect linear relationship (R^2^ of the ‘stretch’ phase of cycles 2 to 10 in **Figure 10 i)** is 0.999 ± 0.00025).

Following repeated cyclic tensile test, the membranes were subjected to 10 minutes of continuous stretching at the rate of 3 mm per minute or until failure, whichever was achieved first. Young’s modulus was calculated and can be described as the ability of the material to store potential energy and release it upon deformation (Schiavi and Prato 2016). The higher the modulus, the stiffer it is. NaOH 1 M 15 minutes treatment group exhibited significantly higher modulus than untreated membranes **(Figure 10.ii.)**. These membranes were also the fastest to fracture during the tensile test. These two findings suggest that this treatment makes the membrane more brittle.

### Permeability assay

Permeability assay was performed as an indirect measure of total membrane permeability. **Figure 11** shows the concentration of FITC that permeated through the membrane at different time-points, expressed as a percentage of the original FITC concentration that was pipetted into the transwell insert (0.0185 mmol). Membranes treated with NaOH 1 M 30 minutes exhibited higher permeability than untreated membranes. This correlates with WCA results as improved hydrophilicity implies more spreading of water droplet across the membrane, thereby increasing the contact area of FITC to the membrane at any given time, facilitating permeation. The percentage of permeated FITC did not increase with time, suggesting that concentration of FITC had equilibrated across the membrane after 10 minutes of permeation.

**Figure 11.**
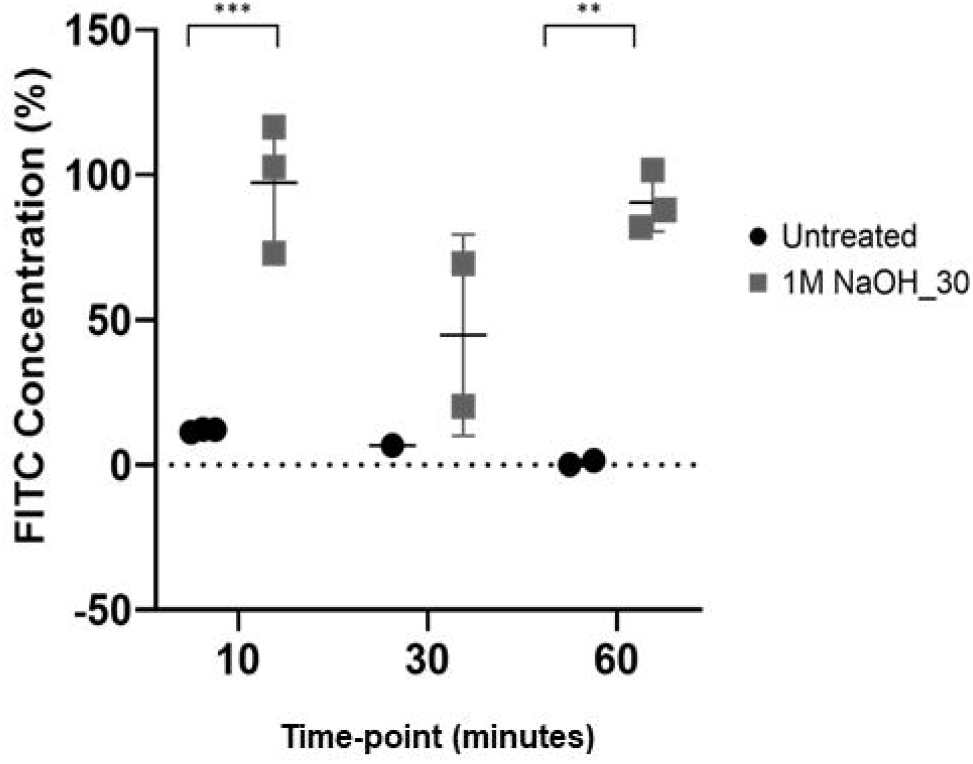
Membranes treated with 1 M NaOH for 30 minutes was more permeable than untreated membrane at the 10- and 60-minute time-point during the assay. Permeability assay was performed on the untreated group and NaOH 1 M 30 minutes group as an indirect measure of total membrane permeability. FITC was used for fluorescence detection. Data are shown as mean ± standard deviation where applicable. Two-way ANOVA: F_(1, 8)_ = 50.6, df = 1, p-value = 0.0001. Dunnett’s multiple comparison: ** p=0.0014, *** p=0.0009

### Overview Discussion

The challenges of developing bioengineered intestinal constructs are complex; not only because of the multiple cell types and layers that interact within the native small intestine, but because the construct must also conform to adequate vascularisation (cultured or engineered) to enable absorption and secretion, as well as withstand/undergo peristaltic action to move digested bulk through the construct, all while maintaining its structure. To tackle this will require a joint-venture between multiple research groups of bioengineers, materials scientists, cell biologists, and clinicians (Clever et al 2019).

Our material has potential to be used in small bowel bioengineering applications; including 3D models or as a potential for TESI. We anticipate further manipulating the material to conform to the tubular structure of the native small bowel to take forward for 3D in vitro cell or organoid culture. Patient et al. have developed a proposed 3D intestinal in vitro model that showed collagen I treated electrospun PET improved Caco-2 cell monolayer formation which exhibited transepithelial electrical resistance properties similar to ex vivo porcine tissue (Patient et al. 2019). Interest has been raised in developing a wholly engineered approach to developing a functional, off-the-shelf product in application for small bowel transplants (Clevers et al. 2019). Our contribution adds to this area of research by presenting a biomaterial that has been reported to be suitable for intestinal cell culture (Patient et al 2019) which can undergo cycles of mechanical stress after exposure to acids, alkali and proteins and is capable of maintaining its structural and physical integrity.

## Conclusion

Our study investigated the use of electrospun PET as a potential biomaterial for small intestine bioengineering applications. We characterised the effects of polymer concentration, collecting distance, voltage, flow rate, collecting time, temperature, humidity, and surface treatment on electrospun PET scaffolds. Polymer concentration played the most significant role in determining fibre morphology and diameter. PET fibres were produced at nano-scaled diameters and formed robust membranes with good mechanical properties. Although electrospun membranes were relatively hydrophobic, the surface chemistry of PET was hydrophilic. Treating membranes with 1 M NaOH for 15 minutes yielded significant improvements in WCA, fibre thinning, rougher membrane topography and enhancement of hydroxyl groups to the polymer surface, with no compromise to the mechanical integrity and membrane elasticity. The material fabricated was porous and permeability properties were improved post-treatment. Electrospinning could, therefore, be used as a potentially cost-effective alternative to fabricate a biomaterial for small bowel engineering applications.

## Author Contributions

KK carried out the data collection and analyses, wrote the entire manuscript and made the figures and tables. AH led the research, contributed to writing the manuscript, reviewed and edited the manuscript.

## Acknowledgements

The authors would like to acknowledge Alison Beckett (University of Liverpool) for her help with the SEM to image the fibrous membrane and Dr Keith Arnold (Materials Innovations Factory, Liverpool) for his help with SEM in imaging the membranes after surface treatments.

