## Supplementary figures and images for "Development of Permanent Artificial Bowel Replacement Substrates"

### supplemental figure 1

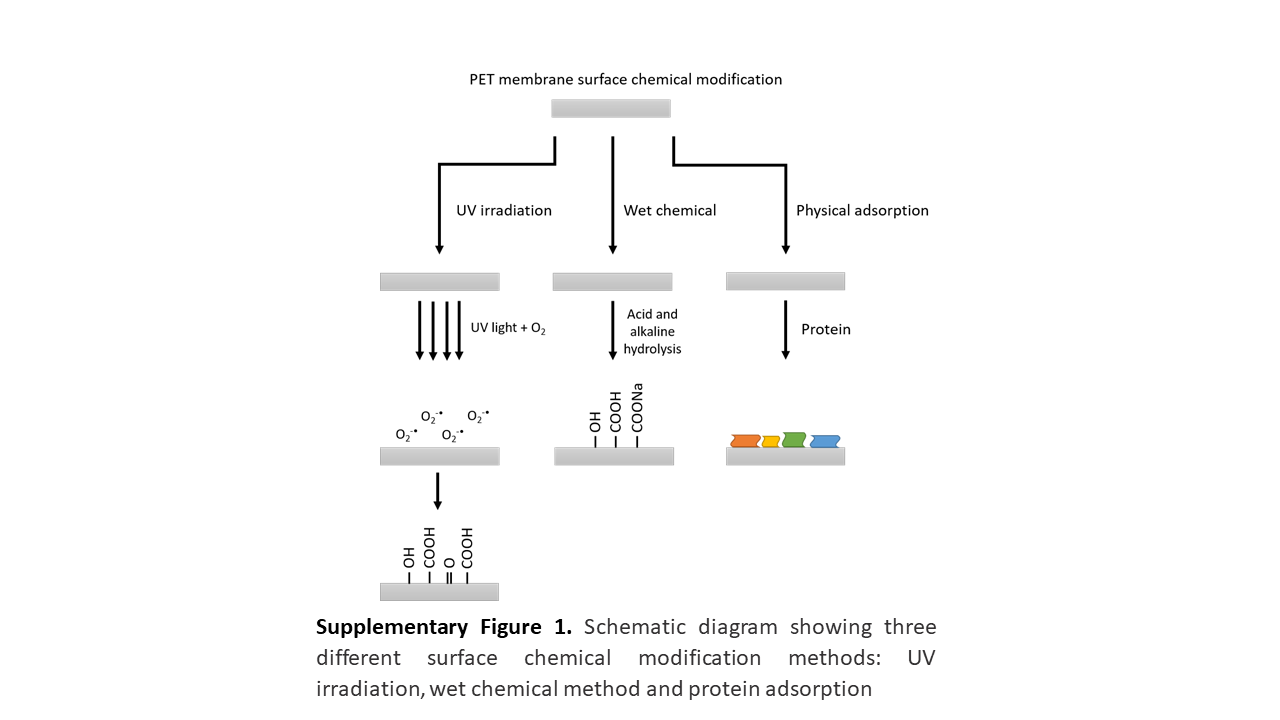

### supplemental figure 2

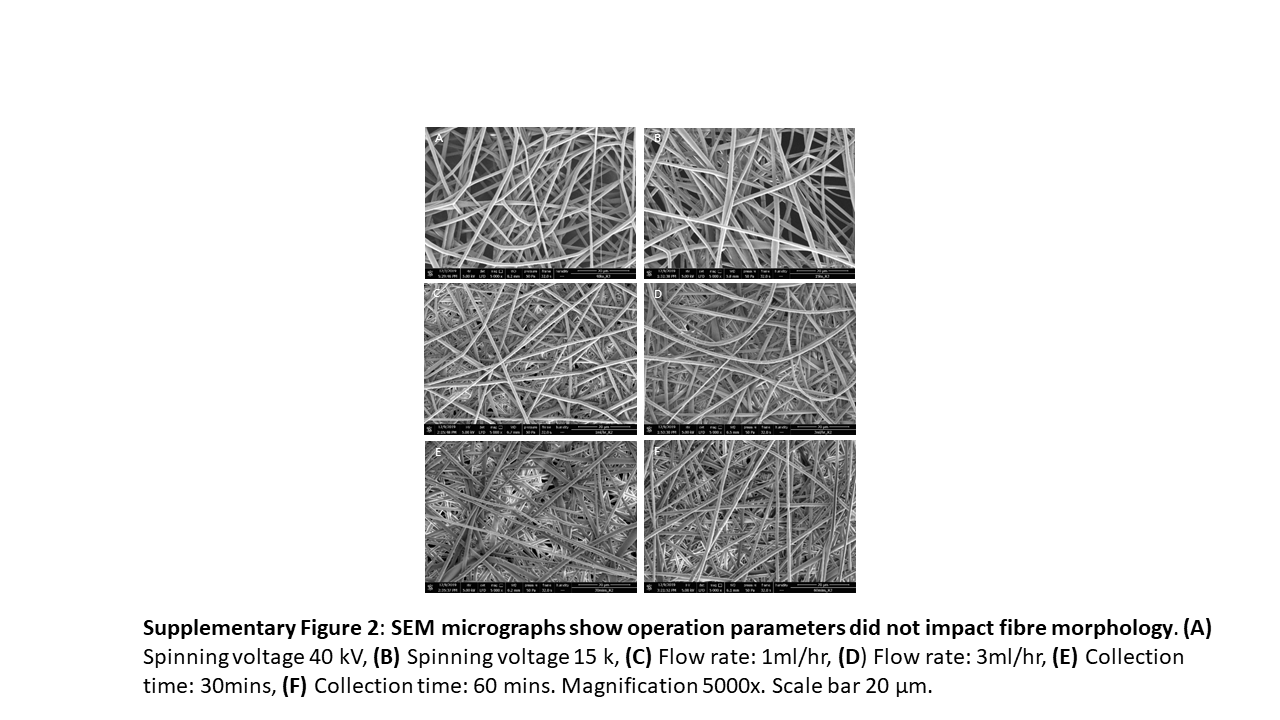
